# Bacterial Volatile Organic Compound Specialists in the Phycosphere

**DOI:** 10.1101/2024.06.14.599129

**Authors:** Vaishnavi G. Padaki, Xavier Mayali, Peter K. Weber, Stephen J. Giovannoni, Kaylene Abraham, Kerry Jacobs, Lindsay Collart, Kimberly H. Halsey

**Affiliations:** Department of Microbiology, Oregon State University; Lawrence Livermore National Laboratory, Livermore, CA 94550 USA

**Keywords:** Phycosphere, Volatile organic compounds, Algal-bacterial interactions, Phytoplankton photosynthesis, Hydrocarbon metabolism

## Abstract

Labile dissolved organic carbon (LDOC) in the oceans accounts for ∼¼ of marine primary production and turns over with a half-life of seconds to days, fueling one of the largest engines of microbial heterotrophic production on the planet. Volatile organic compounds (VOCs) are poorly constrained components of LDOC. Here, we detected 78 *m*/*z* signals, corresponding to unique VOCs, including petroleum hydrocarbons, totaling 18.5 nM in the culture medium of a model diatom. In five cocultures with bacteria adapted to grow with this diatom, 1 to 66 VOCs were depleted. Two of the most active VOC consumers, *Marinobacter* and *Roseibium,* contained more genes encoding VOC oxidation proteins, and attached to the diatom, suggesting VOC specialism. With NanoSIMS and stable isotope labeling, we confirmed that *Marinobacter* incorporated carbon from benzene, one of the depleted VOCs detected in the co-culture. Diatom gross carbon production increased by up to 29% in the presence of VOC consumers, indicating that VOC consumption by heterotrophic bacteria in the phycosphere – a region of rapid organic carbon oxidation that surrounds phytoplankton cells – could impact global rates of gross primary production.

## Introduction

The majority of algal primary production in the world oceans is labile dissolved organic carbon (LDOC), which is readily catabolized by heterotrophic bacteria ^1^. Volatile organic compounds (VOCs) are an important subset of the LDOC produced by phytoplankton ^2^, but VOCs are often overlooked in metabolomic studies because most are not captured by common methods of analysis such as solid phase extraction and liquid chromatography. VOCs are involved in complex allelochemical processes in the marine ecosystem ^3^. Some VOCs defend against algal predation (reviewed in Saha and Fink 2022) ^3^, but dimethylsulfide (DMS) stimulates grazer activity ^4^. VOCs can inhibit algal competitors ^5^ and signal within-population stress ^6^. VOC emissions from the ocean into the atmosphere contribute to secondary aerosols and alter the oxidative capability of the atmosphere. These climate impacts make understanding the biological processes that control VOC accumulation in the surface oceans of utmost importance.

Some VOCs, such as isoprene and DMS, are produced by phytoplankton in response to stress ^4,7,8^, while others (e.g., acetaldehyde and methyl iodide) are intermediate chemicals in metabolic pathways ^9,10^ that can diffuse across cell membranes due to their low molecular weights and hydrophobicity. These diffusive properties may allow VOCs to escape biochemical transformation and cross the algal cell membrane. Other VOCs, including aromatic hydrocarbons such as benzene, toluene, and ethylbenzene/xylene (collectively known as BTEX), are produced by phytoplankton via the shikimate and non-mevalonate pathways. ^11,12^

The multifaceted effects of VOCs suggest they have a role in structuring the microbial community. Research on VOC transfer between algae and bacteria shows that VOCs, including DMS, released by algae can be growth substrates for marine bacteria^13,14^. For example, a wide range of VOCs were detected during the growth of the diatom *Thalassiosira pseudonana*, and a subset of those supported the growth of *Pelagibacter ubique* HTCC1062, a member of the ubiquitous SAR11 bacterioplankton ^10,15^. These publications suggested that VOCs were “public goods” released by the diatom constitutively and available to any organism able to metabolize them. *P. ubique* metabolized isoprene, acetone, acetaldehyde, DMS, and cyclohexanol, which include some of the most well-studied VOCs for their roles in air-sea exchange and atmospheric chemistry ^15^. BTEX are produced by some algae ^11^, and are growth substrates for a wide variety of bacteria isolated from oilfields ^16,17^, soils ^18^, groundwater ^19^, and wastewater ^20^. Less is known about BTEX consumption by marine bacteria, which is likely an important sink that limits BTEX emissions into the atmosphere ^2^. Genes encoding proteins that oxidize some VOCs (e.g., methanol, DMS, acetone, acetaldehyde) in bacteria have been elucidated ^14^. In the ocean, the widespread expression of some of the genes involved, such as acetone/cyclohexanone monooxygenase, suggests marine bacteria have an important role in mediating VOC air-sea emissions ^10^.

The phycosphere, the layer of water immediately surrounding a microalgal cell, is the site of complex interactions between phytoplankton and bacterioplankton, which influence carbon and nutrient cycling in marine ecosystems ^21–23^. Microbial activities in the phycosphere are believed to be distinct from processes occurring in bulk water, a result of bacterial taxa with specialized adaptations populating the region of concentrated chemicals surrounding phytoplankton ^24,25^. Direct chemical transfer from phytoplankton to bacterioplankton should be enhanced by their occupation of the narrow zone of the phycosphere, which allows access to metabolites before they diffuse into the bulk seawater ^22,26^. The ecology of the phycosphere is similar to its terrestrial analog, the rhizosphere: phylogenetically related bacterial taxa are found across both systems ^27^, chemotaxis plays a central role in accessing the narrow zone of high chemical concentration ^28,29^, and the chemicals transferred from the primary and secondary producers are functionally similar ^22^. For example, primary metabolites, such as sugars, amino acids, and vitamins, as well as metabolite precursors, such as organosulfur compounds (dimethylsulfide (DMS) and dimethylsulfoniopropionate) ^30,31^, are used as bacterial growth substrates in both the phycosphere and rhizosphere ^32,33^.

The bacterial community in the phycosphere of marine microalgae may be uniquely adapted to VOC metabolism, as is the case for methylotrophs in the phyllosphere ^34^. The phycosphere of a modest-sized (∼20 μm) diatom extends about 50 – 2000 μm away from the cell ^22^. In this region, chemicals released by algae are modeled to be ≥50% higher in concentration than in bulk seawater ^22^, where VOC concentrations are in the fM to nM range. VOC concentration in the phycosphere is partly controlled by the rate of diffusion from the algal source. The rate of diffusion decreases with increased molecular size and decreased hydrophobicity and likely increases with phytoplankton growth rate and bacterial VOC uptake. Some bacteria occupy the phycosphere through chemotaxis or physical attachment to algal cells ^35,36^. Chemotactic bacteria experience transient exposures to a dynamic organic chemical cocktail in the phycosphere ^32^, where the composition depends on the algal producer ^21,37^, environment ^38,39^, and antagonistic and mutualistic interactions ^32,40,41^, requiring bacterial consumers to efficiently shift physiologies to match resource availability. Strategies bacteria use to navigate carbon acquisition as they traverse phycosphere boundaries are not known but may depend on pool sizes and the diversity of regulatory behavior systems for the uptake and metabolism of various substrates ^42^. Bacteria that can adhere to algae would, in principle, optimize diffusion-mediated VOC uptake by positioning themselves most closely to the algal source.

Diatoms can develop massive spring blooms and are responsible for about half of photosynthesis in the oceans ^43^. *Phaeodactylum tricornutum* is a model diatom whose photophysiology has been extensively studied for ecological and bioenergy purposes. Here, we investigated VOC transfer from the model diatom, *Phaeodactylum tricornutum,* to phycosphere bacteria previously isolated from an outdoor *P. tricornutum* production pond. The bacteria varied in taxonomy, DOC uptake, and attachment to the diatom, and are representive of families (*Rhodobacteriacea*, *Alteromonadaceae*) ubiquitously found in biofuel production ponds as well as the oceans ^32,44,45^. Results show VOC transfer from *P. tricornutum* to its phycosphere bacteria fuels primary and secondary production. Extrapolated to larger scales, our findings support the perspective that phycosphere bacteria are a sink for up to 29% of gross carbon production through VOC transfer, and they play a critical role in limiting VOC accumulation in the surface ocean.

## Materials and Methods

### Culture growth and <u>P. tricornutum</u> physiology

Phaeodactylum tricornutum strain CCMP 2561 was maintained axenically and in separate co-cultures with Roseibium sp. 13C1, Yoonia sp. 4BL, Marinobacter sp. 3-2, Rhodophyticola sp. 6CLA, and Stappia sp. ARW1T (herein “PT-bacteria genus name”). Cultures were grown under 12h:12h light: dark cycles (60-70 µmol photons m^-2^ s^-1^), at 19°C in f/2+Si medium (ASW) ^46^. DAPI (NucBlue, Thermo Fischer) and fluorescence microscopy were used to check P. tricornutum axenicity. Bacterial and diatom growth were determined in triplicate. Cell densities were measured using a GUAVA flow cytometer (Millipore; Billerica, MA, USA) 1-2 h into the light phase. VOCs, chlorophyll content (Chla), and photosynthetic efficiency (F_v_/F_m_) were measured 4-6 h into the light phase. Chla was determined from filtering 2-5 ml of culture (GF/F, Whatman, 25 mm), extracting in 5 ml 90% acetone, and storing at -20° C for 24 h. Chla was quantified by spectrophotometer (Shimadzu; Kyoto, Japan) ^47^. F_v_/F_m_ was measured by fast repetition-rate fluorometer following 10 min dark acclimation ^48^. Short-term carbon fixation rates (approximating gross carbon production, GCP) were measured as in Moore et al. (2020)^15^. In that work, uptake of VOCs by P. ubique HTCC1062 in coculture with T. pseudonana stimulated GCP ∼20% compared to the T. pseudonana monoculture. VOC uptake was confirmed as the cause of the boost in GCP when GCP increased ∼20% after a hydrocarbon trap was installed in the T. pseudonana monoculture to simulate a highly efficient VOC “sink.” In our experiment, axenic P. tricornutum was grown in a closed system with culture headspace recirculating at 80 ml min^-1^ by peristaltic pump and either directed back into the culture medium or passed first through a hydrocarbon trap (Marineland Black Diamond Activated Carbon) and then into the medium. Headspace in P. tricornutum-bacteria cocultures was recirculated in the same manner with no hydrocarbon trap. Flasks containing only ASW were recirculated for 24 h prior to inoculation with the cultures to remove background VOCs. Samples for ^14^CO_2_-uptake (10 ml) were collected by syringe during the P. tricornutum exponential phase (2-4 x 10^5^ cells ml^-1^), spiked with 2 μCi of ^14^C-sodium bicarbonate and incubated 20 min in light or dark. All samples were acidified and vented for 24 h prior to measurement by scintillation counter.

### VOC measurement by PTR-TOF/MS

VOCs were measured in axenic *P. tricornutum* and cocultures by PTR-TOF/MS (PTR-TOF1000, Ionicon, Austria), as in Moore et al. (2020)^15^. VOCs in ASW and HPLC-grade water were measured alongside the samples as blank controls. Culture, ASW, or HPLC water (100 ml) was transferred to a 0.2 L polycarbonate dynamic stripping chamber^49^ and bubbled with breathing-grade air passed first through a hydrocarbon trap and then through a glass frit in the bottom of the chamber at 50 ml min^-1^ for 5 min to strip VOCs from the sample. Stripped VOCs were directed into the PTR-TOF/MS for detection following soft ionization with H_3_O^+^. Mass spectra (30-240 a.m.u.) were acquired every 5 s. Each peak in the spectra is a compound of its molar mass + 1.008 (mass of H^+^). Data from 0.5-5 min were analyzed for VOCs. PTR-TOF/MS data were processed using PTR-viewer 3.4.3 (Ionicon Analytik). h5 files were mass-calibrated using three chemicals known within all spectra (*m*/*z* 29.998, 203.943, and 330.848). First, *m*/*z* signals were binned at 0.5 mass units bounded by 0.25 and 0.75 mass units. Examination of each bin was done to detect multiple peaks within the same bin. A Gaussian-based approach was used to determine the *m*/*z* of each individual peak. Known contaminants, internal standards, water clusters, and fragments of identified parent compounds were removed from further analysis.^52^ Isomeric compounds are not discriminated by PTR-TOF/MS. When possible, *m*/*z* signals were associated with a chemical formula and tentatively identified based on the PTR-Viewer 3.4.3, Ionicon integrated “ChemSpider” database, the GLOVOCS compound assignment database ^50^, and the literature. *m*/*z* signals were categorized into functional groups using the “ChemSpider” database. *m*/*z* signals having isomers in different functional groups were categorized as “mixed.” Integrated peak signals were normalized to H_3_O concentration and detected analyte concentrations (ppbv) calculated without calibration using the simple-reaction-kinetics approach with 30% accuracy for *E/N* values >100Td ^51^ (*E/N* for this study = 126Td).

Analyte concentrations were converted from ppbv to molarity using the formula

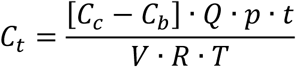

where *C_t_* is the *m*/*z* molar concentration in the *P. tricornutum* monoculture or PT-bacteria coculture, *C_c_ is* the mixing ratio of the *m*/*z* in the headspace of the bubbled culture, *C_b_* is the mixing ratio of the *m*/*z* in the headspace of bubbled ASW, *Q* is the bubbling rate (0.3 L h^-1^), *p* is 1 atm, *t* is the duration of the collected data (4.5 min), *V* is the volume of the culture (0.1 L), *R* is the gas constant (0.0821 L atm K^-1^ mol^-1^), and *T* is temperature (K).

### Benzene uptake and nanoSIMS analyses

Dose experiments were performed to evaluate the growth responses of *Marinobacter, Roseibium, Rhodophyticola, and Stappia* to benzene addition (Supplementary Fig. S1). A saturated benzene solution was prepared in ASW and added to 0.15 L ASW or *P. tricornutum*-spent media (‘PT-spent’ prepared from 0.2 µm filtered spent media collected from axenic *P. tricornutum* grown to 2-4 x 10^5^ cells ml^-1^ and diluted in 1:10 with ASW) in Teflon sealed 0.16 L crimp bottles to obtain final concentrations ranging from 0 to 180 µM prior to bacterial inoculation. Cultures, including no bacteria controls, were incubated at 19°C in the dark and cell density measured daily by withdrawing 0.5 ml by syringe.

^13^C-benzene uptake was evaluated using nanoSIMS in six different treatments grown in Teflon-sealed bottles (*Marinobacter*) or airtight Nalgene bottles (*Rhodophyticola*). 50 nM 98% ^15^N leucine (Cambridge Isotopes, MA, USA) was added to all treatments as an independent measure of cell activity. Sample preparation for nanoSIMS and description of isotopic measurements are in Supplementary Fig. S2.

### SEM imaging

Samples (5 ml) of *Marinobacter, Roseibium,* and *Rhodophyticola* cocultured with *P. tricornutum* were collected in mid-exponential phase, treated with 1.25 ml of 2X 5% glutaraldehyde + 2% paraformaldehyde (PFA) in 0.1 M sodium cocadylate buffer (NaCB), incubated for 24 h, filtered on a 0.2 µm polycarbonate filter, and washed three times with NaCB + Milli-Q water. The filtered cells were subjected to critical point drying and then vacuum-dried and gold-coated. Imaging was done using a detector with a 5.1 mm working distance, 1.8 kV voltage, and 20-30 µm aperture diameter at the Electron Microscope facility, LPSC, Oregon State University. The number of bacteria adhered to *P. tricornutum* in coculture was estimated from 50 diatom cells (Supplementary Table S1).

### Bacterial genes encoding hydrocarbon metabolism

Whole genome sequences of each bacteria were acquired from NCBI (GB accession numbers: *Marinobacter*: PRJNA500125; *Rhodophyticola*: PRJNA441682; *Yoonia*: PRJNA441685; *Stappia*: PRJNA441689; *Roseibium*: PRJNA441686). Geneious Prime v2023.2.1 and Hidden Markov Models were used to search genomes for genes encoding proteins initiating hydrocarbon oxidation (i.e., *alkB*, Pfam(PF00487); *almA*, Pfam(PF00743); *rhdA*, Pfam(PF00848); *acmA*, Pfam(PF00743); *aldH*, Pfam(PF00171) (Supplementary Table S2) ^52^. Each hit for *alkB* and *rhdA* was manually curated using Geneious Prime v2023.2.1. Each hit with bit-score of ≥50 and e-value > 0.001 ^53^ was aligned using MUSCLE protein alignment ^54^ in Geneious prime v2023.2.1.

### Statistics

Statistical analyses and figure construction were done in R studio v.4.1.1, with ggplot2 and Complex heatmap packages ^55,56^. Paired student t-tests (p ≤ 0.05) were used to identify *m*/*z* signals that were detected at different concentrations between treatments. The Benjamini-Hochberg procedure was used to control the false detection rate with *m*/*z* signals with Q-values > 0.1 removed from further analysis. Principle component analysis and Permutational Multivariate Analysis of Variance (PERMANOVA) were used to evaluate VOC accumulation and depletion patterns between cultures.

## Results

### <u>Phaeodactylum tricornutum</u> produced a wide range of VOCs in culture

During exponential growth of *P. tricornutum*, 78 *m*/*z* signals were detected by proton transfer reaction mass spectrometry (PTR-MS) at concentrations higher than the media control [multiple t-test corrected p-value at 0.1 (FDR_B-H_<0.1, n = 6)]. VOCs accumulated in the culture to a total of 85.1 fmol cell^-1^, or 18.5 nM, and represented eight known chemical functional groups (Fig. 1). Hydrocarbons represented about 60% of the VOCs produced in the exponential phase (54.2 fmol cell^-1^; Supplementary Table S3) and included BTEX compounds (*m*/*z* 79.11, *m*/*z* 93.07, and *m*/*z* 107.08), respectively, previously reported in *P. tricornutum* ^11^, and C_11_H_16_ (*m*/*z* 149.13). Alcohols were the second most abundantly produced group of VOCs, contributing up to 11% of the total VOC pool (10.2 fmol cell^-1^). Methanol (*m*/*z* 33.03) and, likely, ethanol (*m*/*z* 47.04) made up the vast majority of the accumulated alcohols. Consistent with reports that *P. tricornutum* is a halocarbon producer ^57,58^, *m*/*z* 124.96, corresponding to bromoethanol, was detected in the exponential phase. The well-studied VOCs acetone (*m*/*z* 59.05), acetaldehyde (*m*/*z* 45.03), and DMS (*m*/*z* 63.03) were also detected in the exponential phase.

**Fig. 1:**
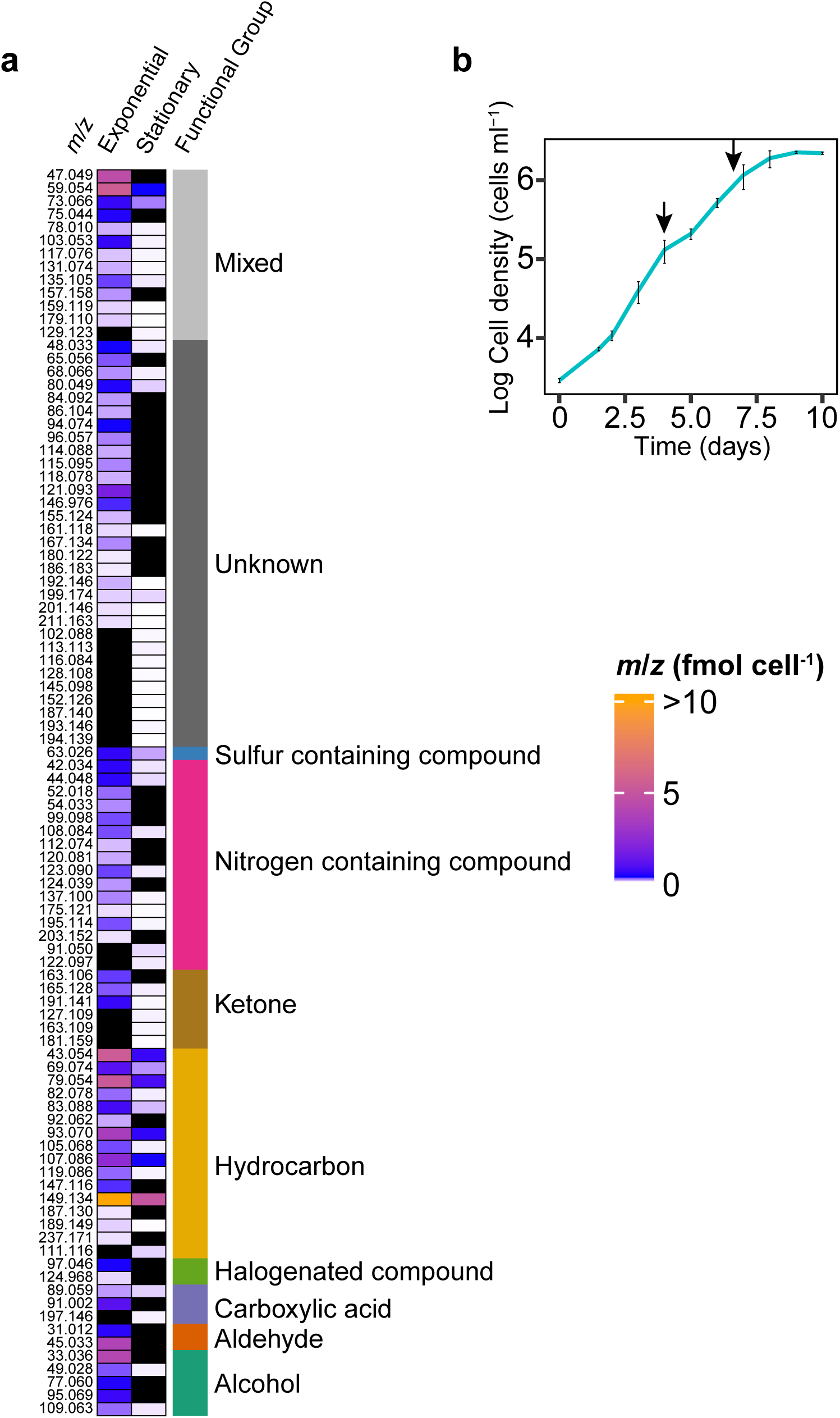
VOC production in *P. tricornutum*. **a** *m*/*z* signals (rows) produced during exponential and stationary phases. The color scale ranges from low (light blue) to high (orange) production. Black are *m*/*z* signals that were not detected in higher amounts in one of the phases than the media control. Functional group categorization of each *m*/*z* signal is in the rightmost column. **b** *P. tricornutum* growth (Error bars are SE for n=6 independent cultures). Arrows indicate sampling days for VOCs.

After *P. tricornutum* entered the stationary growth phase, 58 *m*/*z* signals were detected in concentrations higher than the media control (FDR_B-H_<0.1, n = 6) and accumulated to 9.01 fmol cell^-1^, or 0.96 fM (Fig. 1), which was only about 10% of the VOC accumulation during the exponential phase (Fig. 1, Supplementary Table S3). VOCs detected in both phases of growth were consistently lower in concentration in the stationary phase. Slower VOC production in response to cessation of growth, and loss of VOCs from the vented growth flasks, are the most likely explanations for the concentration differences between growth phases. Hydrocarbons still represented most of the VOCs (84%) but their production was only 7.59 fmol cell^-1^. Sixteen of the 58 *m*/*z* signals were unique to the stationary phase, but many of these had unknown identities.

The most abundant VOC in both exponential and stationary phases was *m*/*z* 149.13, accumulating to 32.5 and 4.9 fmol cell^-1^ in each growth phase, respectively. This *m*/*z* signal corresponds to the unprotonated formula C_11_H_16_, a hydrocarbon with isomers including dictyopterene D’, which was previously detected in brown algae^59,60^. *m*/*z* 149.13 also corresponds to the aromatic hydrocarbons, neopentyl-benzene, 4-tert-butyl-toluene, and 1-ethyl-2-propylbenzene, which have been identified in plant leaves ^61^. Forty-two *m*/*z* signals, including BTEX, were detected in both exponential and stationary phases (Fig. 1).

### VOCs were depleted in <u>P. tricornutum</u>-bacteria cocultures

Five *P. tricornutum*-bacteria cocultures were grown to identify VOCs produced by the diatom that decreased in concentration in the presence of a bacterium, which would suggest those VOCs were consumed by the bacterium. Alternatively, lower VOC concentrations in a coculture could indicate VOC production was inhibited in the presence of the bacterium. The five *P. tricornutum*-bacteria cocultures (herein “PT-bacteria genus name”) each showed distinct VOC depletion patterns during exponential growth when compared to axenic *P. tricornutum* (Fig. 2a, Supplementary Fig. S3, PERMANOVA, p-value = 0.03, n = 6). Of the 78 *m*/*z* signals produced by *P. tricornutum* in exponential phase, 31 to 59 were significantly depleted in four of the five cocultures (FDR_B-H_<0.1, n = 6) (Fig. 2a). VOCs were most strongly depleted in PT-*Marinobacter* (7.09 fmol cell^-1^), followed by PT-*Stappia* (6.51 fmol cell^-1^), PT-*Roseibium* (5.52 fmol cell^-1^) and PT-*Rhodophyticola* (4.54 fmol cell^-1^), where VOC depletion was normalized to the concentration of bacterial cells in the coculture (Fig. 2a, Fig. 4, Supplementary Table S4).

**Fig. 2:**
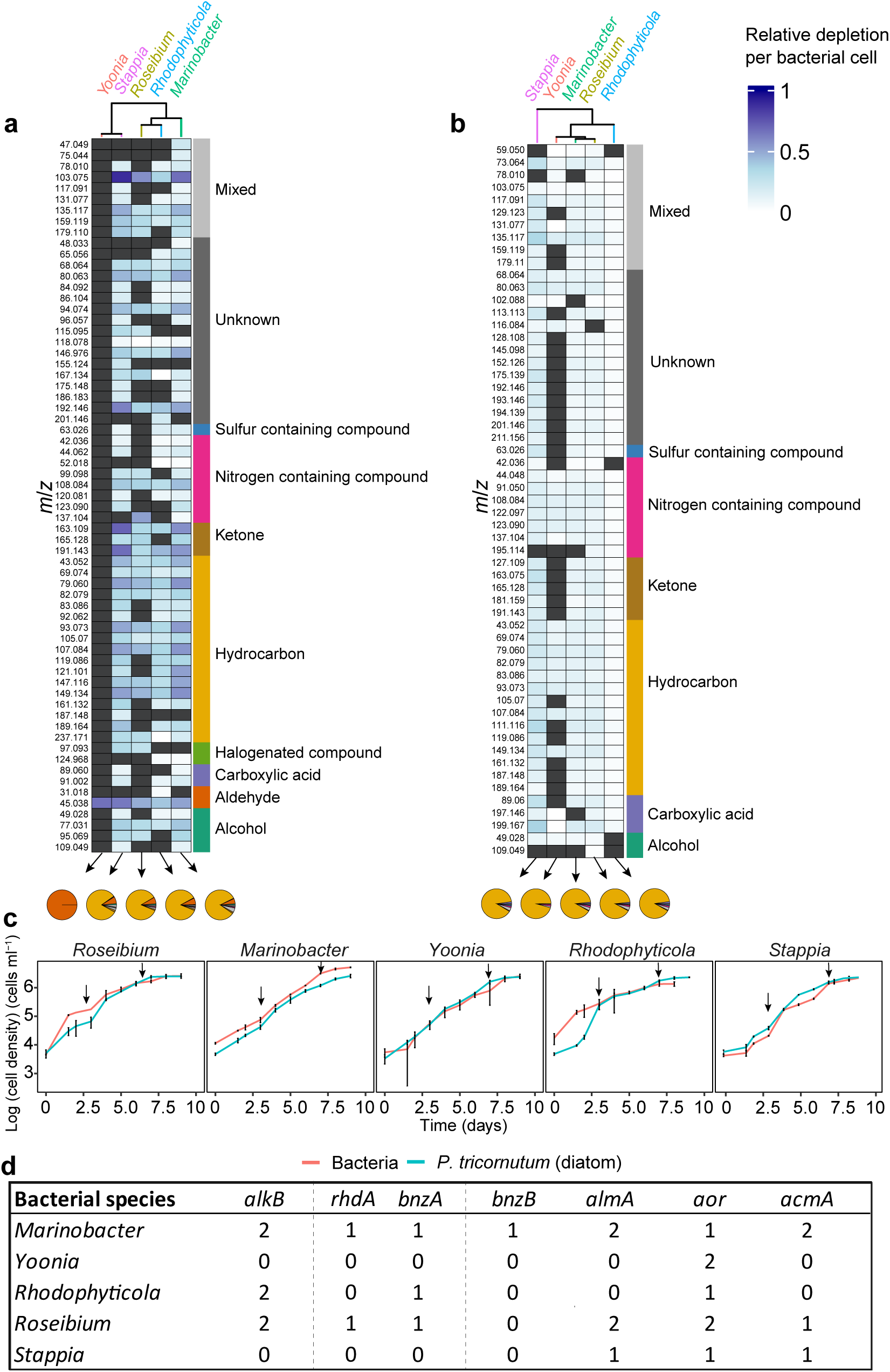
Phycosphere bacteria demonstrate widely ranging VOC depletion patterns. **a,b** *m*/*z* depletion (rows) in each *P. tricornutum*–bacteria coculture (columns are labeled with the bacteria grown with *P. tricornutum*) in exponential (**a**) and stationary phase (**b**) relative to axenic *P. tricornutum* in each of the growth phases. The magnitude of depletion is shown, ranging from low (light blue) to high (dark blue). Charcoal grey are *m*/*z* signals that were not depleted in cocultures relative to the *P. tricornutum* monoculture. Pearson correlation distances were used to cluster depletion patterns between PT-bacteria cocultures (top axes). Pie charts show the proportion of each functional group depleted. **c** Growth of each bacteria (red lines) in coculture with *P. tricornutum* (blue lines), (Error bars are SE, n=6). Arrows indicate sampling days for VOCs. **d** Genes encoding hydrocarbon metabolism proteins in the phycosphere bacteria. Number of gene copies is shown for each bacterium (bit-score ≥50, e-value >0.001). Abbreviations: *alkB* (alkane-1-monooxygenase), *bnzA* (benzene dioxygenase alpha-subunit), *bnzB* (benzene dioxygenase beta-subunit), *almA* (flavin-binding monooxygenase), *aor* (aldehyde oxidoreductase), *acmA* (acetone monooxygenase).

About half the number of *m*/*z* signals were depleted in the PT*-Roseibium* compared to PT-*Marinobacter*. Some *Marinobacter* and *Roseibium* cells were observed to be attached to *P. tricornutum* (Fig. 3) ^44^, thus the cell-normalized VOC depletion data in PT-*Marinobacter* and PT-*Roseibium* are based on estimates of bacteria cell densities using flow cytometry for the free-living and scanning electron microscopy for the attached bacteria (see Methods, Supplementary Table S1). In PT-*Yoonia*, only one *m*/*z* signal, 45.03, corresponding to acetaldehyde, was significantly depleted (0.5 fmol cell^-1^ (Fig. 2a, Supplementary Table S4). There were 14 *m*/*z* signals that did not show significant depletion in any of the five PT-bacteria cocultures. A lack of depletion could mean that the bacteria could not incorporate or metabolize those VOCs, or those VOCs were maintained at a steady state concentration by equal rates of production and consumption in the coculture. Of those, 5 *m*/*z* signals belonged to the nitrogen-containing category of VOCs (Supplementary Table S5). Furthermore, neither methanol nor acetone were depleted in any of the five cocultures.

**Fig. 3.**
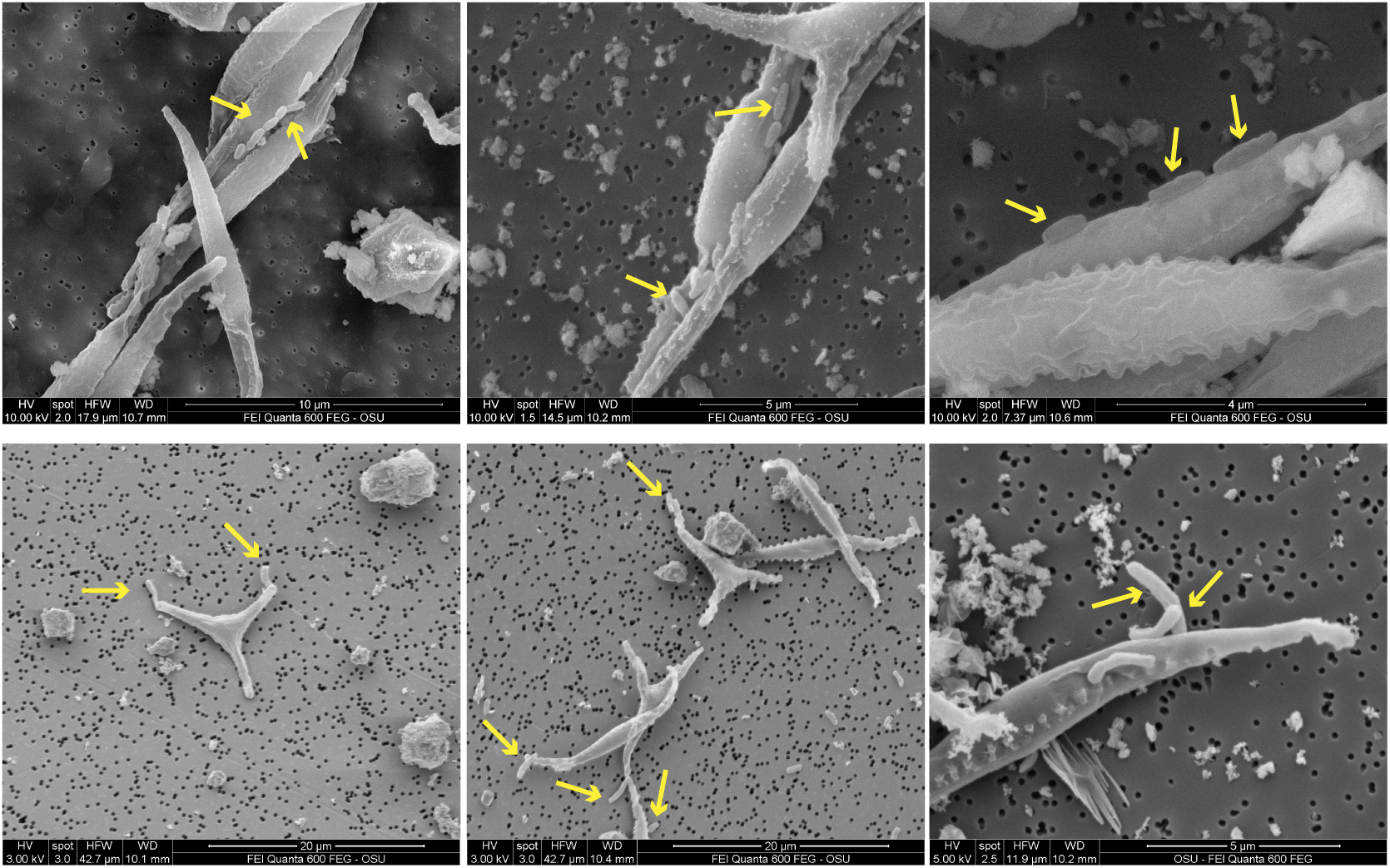
Scanning electron microscopy images showing (top panels) *Marinobacter* attached to *P. tricornutum* in a flat orientation and (bottom panels) *Roseibium* attached to *P. tricornutum* in a polar orientation.

Hydrocarbons were the most highly depleted group of VOCs in all PT-bacteria cocultures except PT-*Yoonia*. Depending on the bacterium present, 10-17 of the depleted *m*/*z* signals were classified as hydrocarbons and made up to 70% of the total VOC pool depleted in the cocultures. PT-*Marinobacter* depleted the most hydrocarbons (5.95 fmol cell^-1^), followed by PT-*Stappia* (5.44 fmol cell ^-1^), PT-*Roseibium* (4.48 fmol cell^-1^) and PT-*Rhodophyticola* (3.94 fmol cell ^-1^) (Fig. 2a). Monoaromatic hydrocarbons, including BTEX, C_11_H_16_, and *m*/*z* 103.05 (likely corresponding to phenylacetylene), were the most strongly depleted hydrocarbons in the PT-bacteria cocultures (Fig. 2a).

In stationary phase, the range of depleted VOCs in the PT-bacteria cocultures was lower, from 0.64-4.97 fmol cell^-1^ (average 2.96 ± 0.66 fmol cell^-1^). The depletion patterns between PT-bacteria cocultures in stationary phase were not distinct from one another (Fig. 2b, Supplementary Fig. S3, PERMANOVA p-value = 0.22, n = 6). In total, 24 to 55 *m*/*z* signals were depleted in the PT-bacteria cocultures (FDR_B-H_<0.1, n = 6). The number of *m*/*z* signals depleted in PT-*Yoonia* expanded from 1 in the exponential phase to 24 in the stationary phase, but the amounts of the individual VOCs depleted were low (Fig. 2b, Supplementary Tables S4 and S7). In stationary phase, 8-14 *m*/*z* signals depleted in PT-bacteria cocultures were classified as hydrocarbons, and these VOCs were the majority of the depleted VOC pools (80-92%). VOCs classified as carboxylic acids were the second-most depleted group, followed by ketones (Fig. 2a and b). Except for hydrocarbons, the magnitude of VOC depletion in PT-bacteria cocultures did not mirror VOC abundances in axenic *P. tricornutum* in either growth phase.

### Bacterial VOC uptake comprised up to 29% of gross carbon production

*P. tricornutum* growth rates, chlorophyll content, and photosynthetic efficiencies were unaffected by the presence of any of the bacterial species in the PT-bacteria cocultures (Supplementary Table S6). To assess the impact of bacterial VOC uptake on *P. tricornutum* primary production, short-term ^14^CO_2_-uptake rates, approximating gross carbon production (GCP) were measured. GCP in PT-bacterial co-cultures were higher in PT-*Marinobacter,* PT-*Stappia,* and PT-*Rhodophyticola* than in axenic *P. tricornutum* (Fig. 4). To confirm that VOC uptake, rather than non-volatile DOC uptake, increased GCP in the cocultures, a hydrocarbon trap was installed that stripped VOCs from the axenic *P. tricornutum* culture, simulating a highly efficient bacterial VOC sink. This treatment caused GCP to increase by 24.9% compared to axenic *P. tricornutum* with no trap (Fig. 4; p = 0.025, n = 6). GCP in PT-*Marinobacter,* PT-*Rhodophyticola,* and PT-*Stappia* were similar to axenic *P. tricornutum* with the hydrocarbon trap and 20.1% to 29.3% higher than in axenic *P. tricornutum* (p-values < 0.05, t-tests, n=3). These differences in GCP corresponded to 0.05 to 0.12 nmol C cell^-1^ h^-1^ transferred from *P. tricornutum* to the bacteria in the form of VOCs (Fig. 4). Consistent with the minimal amount of VOC depleted in PT-*Yoonia*, there was no difference in GCP between PT-*Yoonia* and axenic *P. tricornutum* (p-value = 0.23, t-test, n=3). GCP was not stimulated in PT-*Roseibium* until *Roseibium* entered the second exponential growth phase in its diauxic growth pattern (Fig. 2; note lag between days 2 and 4). This result suggests that *Roseibium* shifted carbon uptake from primarily LDOC early in culture growth to primarily VOCs later in culture growth. VOC depletion in the PT-bacteria cocultures appears to be low compared to the rate of VOC uptake determined by differences in GCP. However, VOC depletion data likely significantly underestimate bacterial VOC uptake because VOC concentrations in the cocultures reflect only slight differences in steady state rates of VOC production and consumption.

**Fig. 4:**
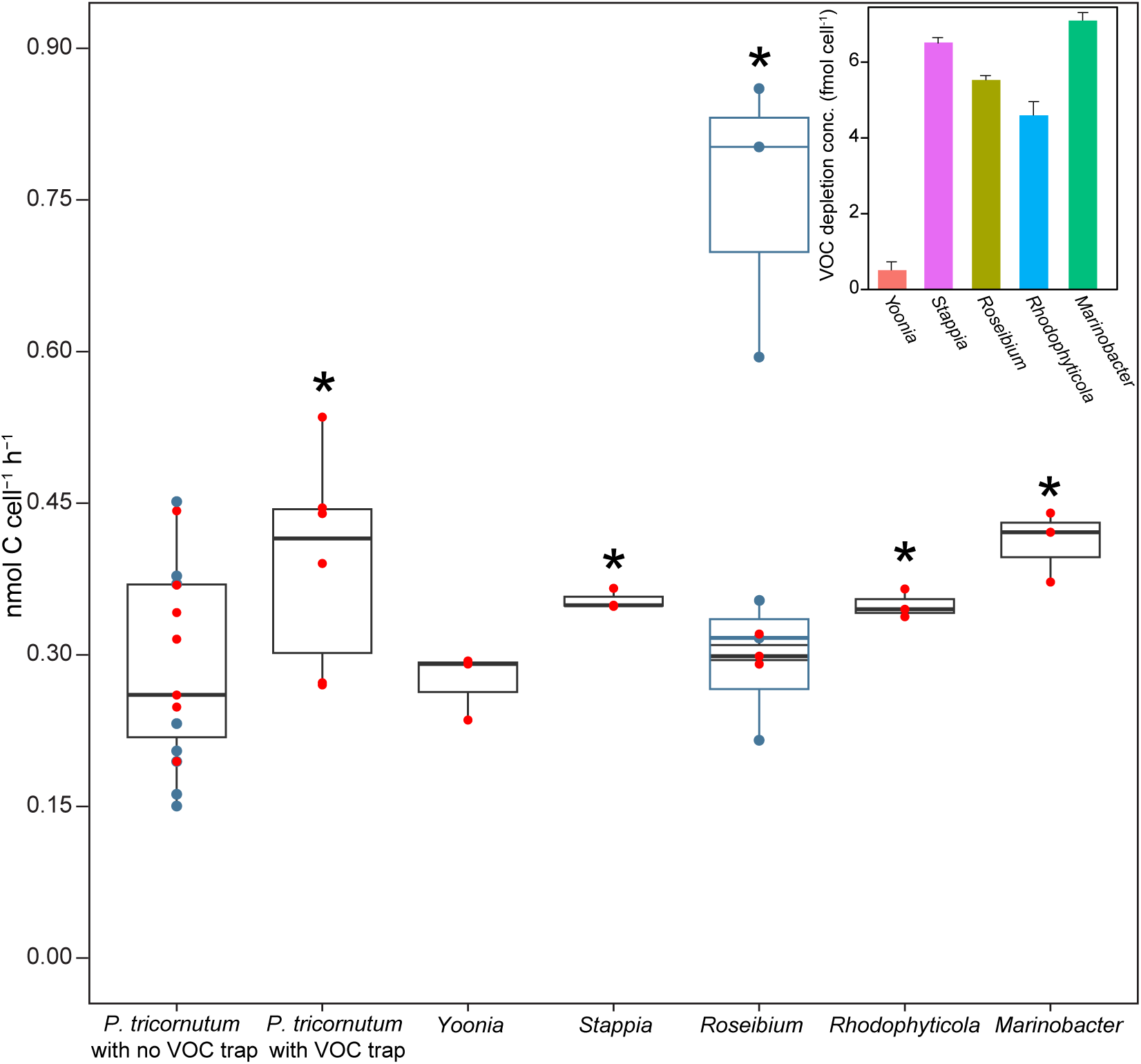
Gross carbon production in *P. tricornutum* in the presence and absence of VOC sinks (hydrocarbon trap or each bacteria in coculture). Points are short-term (20 min) carbon fixation rates, approximating gross carbon production (GCP) for each culture or coculture. Box plots show the mean and upper and lower quartile divisions; red dots show data collected on day 4 (marked in Fig. 2c); blue dots & boxes show data collected on days (1, low box plot, and 5, high box plot) in PT-*Roseibium*); whiskers show 95% confidence limits. Asterisks are GCP in treatments (*P. tricornutum* with VOC trap and PT-cocultures) that were greater than GCP in axenic *P. tricornutum* with no VOC trap for the same day in the growth curve (p-value < 0.05; t-test). Inset shows the bacterial cell-normalized VOC depletion in each *P. tricornutum*-bacteria coculture (Error bars are SE, n = 6).

### VOC depletion correlates with genome content for hydrocarbon metabolism

Of the five bacteria tested*, Marinobacter* and *Roseibium* genomes harbor the largest number of genes encoding hydrocarbon oxidation proteins. For example, genes identified in *Marinobacter* and *Roseibium* included *rhdA,* encoding ring hydroxylating dehydrogenase, *almA*, encoding flavin-binding monooxygenase involved in long-chain alkane degradation, and *alkB,* encoding alkane monooxygenase. RhdA is in a superclass of nonheme iron enzymes that convert aromatic hydrocarbons to dihydrodiols in the presence of dioxygen and NADH ^62^. The broad substrate range for RhdA includes benzene, and the benzene-alkyl substituted hydrocarbons toluene, ethylbenzene, and xylene, consistent with BTEX depletion profiles in *Marinobacter* and *Roseibium*. Benzene dioxygenases are a subclass of RhdA that oxidize benzene and toluene to their corresponding dihydrodiols. We manually curated the *rhdA* hits for *bnzA*, encoding the α-subunit of the benzene 1,2 dioxygenase hydroxylase, based on conservation of the active site motif Cys-X_1_-His-15-to-17 aa-Cys-X_2_-His (where X is any amino acid) ^63^. The gene, *bnzA,* was identified in *Marinobacter*, *Rhodophyticola*, and *Roseibium* (Fig. 2d). The gene, *bnzB,* encoding the β-subunit of benzene 1,2 dioxygenase hydroxylase, was also identified in *Marinobacter* (Fig. 2d). The presence of both α and β-subunits increases hydroxylation activity compared to the activity of either of the subunits alone ^64^. Genes encoding hydrocarbon oxidation in *Stappia* were not identified. Acetone/cyclohexanone monooxygenase, encoded by *acmA* was identified in *Marinobacter, Roseibium, and Stappia*. The broad substrate range of acetone/cyclohexanone monooxygenase ^65,66^ may include cyclic ketones, such as valerophenone (*m*/*z* 163.11) and E-jasmone (*m*/*z* 165.13), corresponding to *m*/*z* signals that were depleted in PT-*Marinobacter*, PT-*Roseibium*, and PT-*Stappia*. One or two copies of *aor,* encoding aldehyde dehydrogenase, were identified in each of the five bacteria, consistent with acetaldehyde (*m*/*z* 45.03) depletion in the five cocultures (Fig. 2).

### Marinobacter and Rhodophyticola exhibited distinct benzene metabolism

We selected benzene as a representative hydrocarbon to directly test incorporation into biomass by *Marinobacter* and *Rhodophyticola*. These bacteria differed in the amounts of benzene depleted in the PT-bacteria cocultures, in their molecular capacities for hydrocarbon metabolism and chemotaxis, in their attachment physiologies (Fig. 2, 3) ^44^, and could be grown without *P. tricornutum* in media prepared from filtrate collected from *P. tricornutum* in exponential growth (‘PTspent’; see methods). Benzene supported growth of *Marinobacter*, but not *Rhodophyticola,* in ASW, and both bacteria grew to higher cell densities in PTspent with added benzene vs. no added benzene (Supplementary Fig. S1; Fig. 5). Benzene uptake and subsequent incorporation into biomass by *Marinobacter* and *Rhodophyticola* were quantified using nanoSIMS. We incubated each bacterium with and without ^13^C benzene (60 µM in *Marinobacter* cultures, 36 µM in *Rhodophyticola* cultures) and ^15^N leucine (50 nM), in f/2+Si artificial seawater medium (ASW), PTspent medium, and in coculture with *P. tricornutum*.

**Fig. 5:**
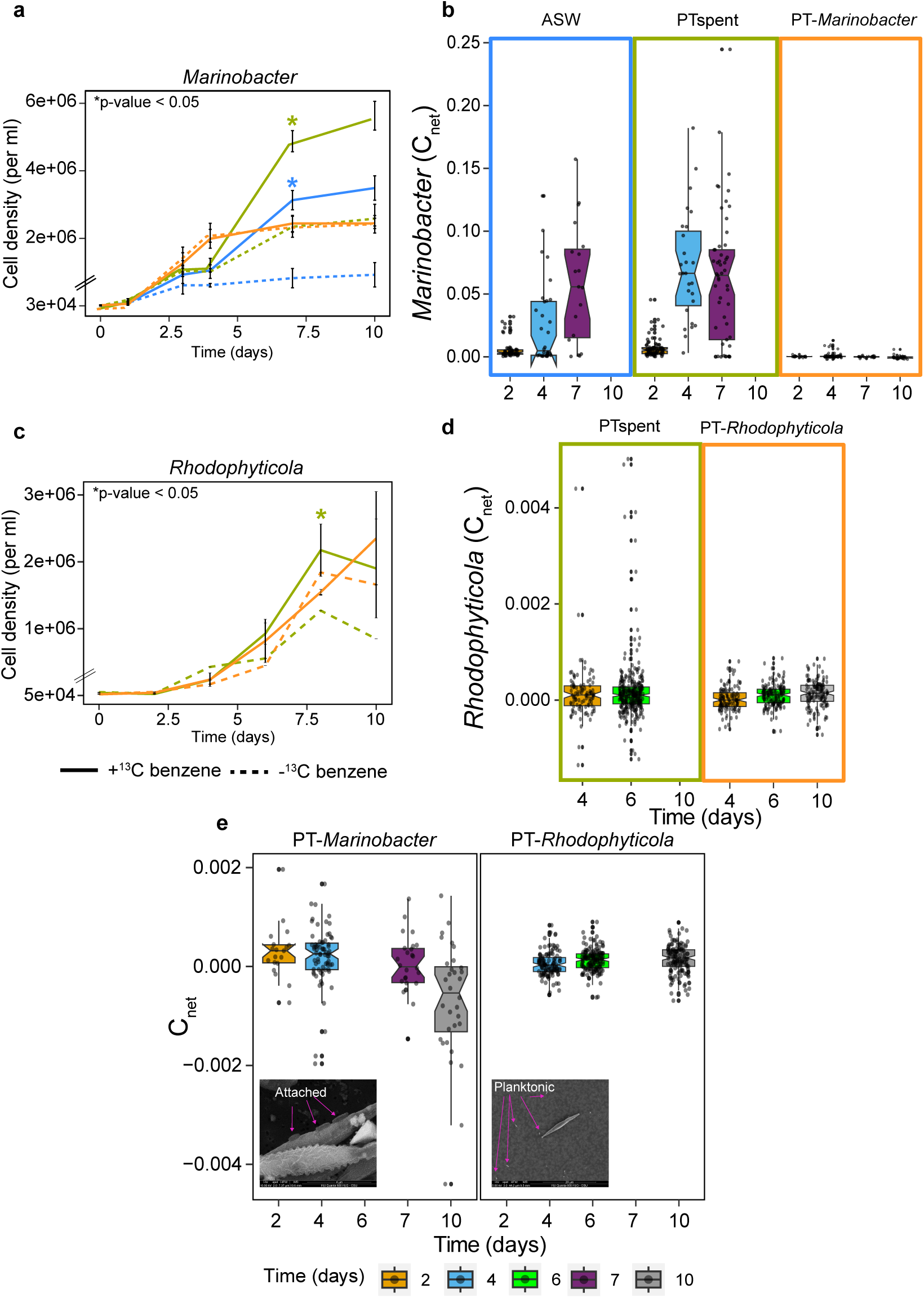
Benzene incorporation into *Marinobacter* and *Rhodophyticola* biomass. **a,c** Bacterial cell densities in the presence (solid lines) and absence (dashed lines) of ^13^C benzene in ASW (blue, not done in *Rhodophyticola*), PTspent (olive), and coculture with *P. tricornutum* (orange). Asterisks indicate treatments with higher cell densities in the presence of benzene compared to the same media treatment with no added benzene (p<0.05, Error bars are SE, n = 3). **b,d** ^13^C enrichment in *Marinobacter* and *Rhodophyticola* in ASW (*Marinobacter* only), PTspent, and in coculture with *P. tricornutum*. C_net_ is the fraction of biomass labeled from ^13^C benzene. **e** Bacterial C_net_ values in PT-*Marinobacter* and PT-*Rhodophyticola* with expanded y-axis. Insets are SEM images of *Marinobacter* attached to *P. tricornutum* and *Rhodophyticola* unattached.

*Marinobacter* cell densities were higher in ASW with benzene added as the sole source of carbon compared to ASW with no benzene added (Fig. 5a p-value <0.05, n = 3). Associated C_net_ values of incorporation, measuring the fraction of a cell’s C originating from benzene, increased over the incubation period, reaching 0.03 to 0.15 (average 0.065, or 6.5%) on day 7 (Fig. 5b).

*Marinobacter* reached higher cell densities in PTspent with added benzene compared to PTspent with no benzene (Fig. 5a p-value <0.05, n = 3). Consistent with benzene-stimulated growth in PTspent, *Marinobacter* C_net_ values in that treatment ranged from 0.02 to 0.17, averaging 0.07 on day 4. *Marinobacter* cell densities were unaffected by benzene addition to PT-*Marinobacter*.

Interestingly, *Marinobacter* did not incorporate ^13^C benzene in the presence of live *P. tricornutum* cells. *Marinobacter* C_net_ values in the coculture averaged zero across the 10-day incubation, even in the presence of 60 µM externally-supplied benzene (Fig. 5e). N_net_ values from leucine in all *Marinobacter* treatments, including the coculture, were low, but higher than killed cell controls (t-tests, p-values <0.01, Supplementary Fig. S2), confirming bacterial activity.

Despite significantly higher *Rhodophyticola* cell densities in PTspent with 36 µM benzene added compared to the no benzene-added control (Fig. 5c), ^13^C-benzene addition to PTspent resulted in low C_net_ values, and only 2% of the population showed enrichment in ^13^C benzene on day 6 (Fig. 5d). *Rhodophyticola* may take up and oxidize benzene for energy rather than biomass generation. No significant difference was observed in *Rhodophyticola* cell densities grown in the PT coculture with or without added ^13^C-benzene. Similar to *Marinobacter*, C_net_ values in *Rhodophyticola* grown in the PT coculture were very low. *Rhodophyticola* showed ^15^N enrichment from leucine in both PTspent and PT-*Rhodophyticola* treatments (Supplementary Fig. S2).

## Discussion

Phycospheres are recognized as energy-rich regions of heterotrophic activity where microbes adapted to this niche benefit from proximity to high concentrations of the organic resources produced by phytoplankton cells. VOCs are diffusible molecules that easily traverse cell membranes and, due to the introduction of new measurement technologies, are increasingly recognized as complex and important products of phytoplankton metabolism. Here, we show that the phycosphere concept applies to VOCs and that they appear to be an important component of photosynthetic carbon transfer to heterotrophic microbial populations in the region immediately surrounding phytoplankton cells. We also show that some phycosphere bacteria have traits that can be interpreted as evidence of VOC specialism, enabling them to access these chemicals whose concentrations are expected to decrease with distance from their source.

VOCs were primarily produced during active *P. tricornutum* growth, when over a quarter of gross carbon production (GCP) was emitted from *P. tricornutum* as VOCs captured in a hydrocarbon trap. Hydrocarbons represented 64% of the total VOCs produced, with BTEX accounting for 13% of the emitted VOC pool, accumulating to 11.8 nM in the diatom culture. BTEX production by phytoplankton has been proposed to be of sufficient magnitude to influence atmospheric chemistry ^11^, unless bacterial VOC consumption prevents BTEX accumulation in the surface ocean ^2^. The VOC concentrations we measured could be underestimates of VOC production because the cultures were grown in vented flasks. Furthermore, in the absence of bacterial sinks, internal VOC concentrations in axenic *P. tricornutum* cells may reach equilibria with the surrounding medium limiting ongoing VOC accumulation in cultures.

We measured concentrations of VOCs in diatom-bacteria cocultures relative to the diatom grown alone to investigate the potential for VOC uptake by bacteria. In the case of benzene, isotopic labeling confirmed that *Marinobacter* used benzene for biomass generation, but for other VOCs, we cannot eliminate the possibility that VOC depletion in coculture was caused by bacterial inhibition of VOC production by *P. tricornutum*. *P. tricornutum* GCP was stimulated in cocultures where depletion of a wide range of VOCs was also observed. Some marine bacteria are known to metabolize VOCs, including BTEX ^15,67^, and we showed that the bacteria that caused VOC depletion also have genes encoding enzymes that oxidize some of the depleted compounds. For other depleted compounds, genetic loci associated with bacterial metabolism are unknown.

The data we report here indicate a range of bacterial VOC specialism (Table 1). Four of the five bacterial strains harbor multiple genes that encode hydrocarbon oxidation enzymes with broad substrate ranges. *Marinobacter* and *Roseibium,* which consumed large amounts of hydrocarbons, had the most genes encoding for hydrocarbon oxidation. *Marinobacter* and *Roseibium* had few genes encoding complex polysaccharide metabolism, and in previous research, incorporated low amounts of macromolecular DOC into biomass ^44^. In our experiments, *Rhodophyticola* and *Stappia,* which harbor fewer hydrocarbon oxidation genes, depleted smaller amounts of individual hydrocarbons but a wide range. Mayali et al. reported that *Stappia, Rhodophyticola,* and *Yoonia* used complex DOC in their 2023 study, in agreement with their larger array of genes encoding oligosaccharide, starch, and cellulose degradation ^44^.

**Table 1:** Traits of bacteria used in this study. Plus symbols indicate the relative use of carbohydrate and VOC metabolism in each bacterium.

| Bacteria | Carbohydrate metabolism<sup>a</sup> | VOC metabolism<sup>b</sup> | Flagellar biosynthesis (*flhA*)<sup>c</sup> | Chemotaxis (*che* genes)<sup>a</sup> | Attachment to diatom<sup>a,b</sup> |
| --- | --- | --- | --- | --- | --- |
| *Marinobacter* | + | +++ | 1 copy | Yes | Yes |
| *Roseibium* | + | +++ | 2 copies | Yes | Yes |
| *Rhodophyticola* | +++ | ++ | Not present | No | No |
| *Stappia* | +++ | ++ | Not present | No | No |
| *Yoonia* | +++ | + | Not present | Yes | Yes |

<sup>a</sup>Summarized from Mayali et al. 2023
<sup>b</sup>Summarized from this study
<sup>c</sup>*flhA* in *Marinobacter* has 72.9% amino acid similarity (e-value 2.58e-158) to *flhA* from *Escherichia coli* (P76298) and *flhA* in *Roseibium* have 64.5% (e-value 5.84e-106) and 67.0% (e-value 4.18e-89) amino acid similarity to P76298..

Thus, for this set of traits, among a set of five heterotrophic isolates from *P. tricornutum* ponds, two strains appeared to be VOC specialists: they used more VOCs and less carbohydrate DOM, and as discussed below, they were motile and attached to *P. tricornutum*. Two strains used lesser amounts of VOCs, are active metabolizers of carbohydrate DOM, are not motile, and did not attach; and one strain used carbohydrate DOM, only a single VOC, and attached.

We searched for evidence of phycosphere occupation to understand whether, in the strains we studied, it was linked to VOC use. The VOC specialists, *Marinobacter* and *Roseibium,* and one strain that did not use VOCs, *Yoonia*, attached to *P. tricornutum,* but the other strains did not. *Marinobacter* attached to the diatom in a flat orientation, while *Roseibium* attached to *P. tricornutum* using polar adhesion (Fig. 3). It has been reported that chemotaxis and motility are key traits of hydrocarbon-degrading bacteria. Volatile aromatic hydrocarbons, including toluene and benzene derivatives, have been shown in previous work to be effective bacterial chemoattractants, even in the presence of complex organic matter, such as petroleum, crude oil, and diesel ^68–70^. *Marinobacter* and *Roseibium* have canonical *che* genes, which encode for chemotaxis ^44^, but chemoattractant compounds have not been identified in these bacteria ^28,71^ (Table 1).

One of the most interesting results in this study was the absence of ^13^C-benzene cellular incorporation in *Marinobacter* grown in coculture with *P. tricornutum,* despite strong ^13^C-benzene incorporation in *Marinobacter* grown in the absence of *P. tricornutum* (i.e., in PTspent and ASW) and significant benzene depletion in PT-*Marinobacter* compared to the *P. tricronutum* monoculture (Fig 5). The experiment was repeated, and the results were confirmed. These results suggest that in coculture with *P. tricornutum, Marinobacter* incorporated ^12^C-benzene, and possibly other hydrocarbons, directly from the diatom and specifically did not incorporate ^13^C-benzene from the bulk medium. *Marinobacter* and *Roseibium* were sometimes observed attached to *P. tricornutum.* Bacterial attachment or entry into the phycosphere ^28^ could facilitate efficient VOC transfer directly from the diatom to the bacteria. In coculture, VOC uptake from the diatom to *Marinobacter* may occur via their physical interaction in flat orientation, enabling passive VOC diffusion across the diatom and bacterial membranes. In addition, FadL outer membrane transporters for hydrophobic molecules can boost BTEX metabolism in some Gram-negative hydrocarbon degraders ^72^, and the *Marinobacter* strain used in this study encodes a FadL protein with 72% nucleotide identity to the well-characterized FadL in *Escherichia coli*. In the presence of the diatom, *Marinobacter* may regulate FadL to optimize use of a wide range of substrates obtained from *P. tricornutum*. However, FadL substrate specificity and kinetics are not yet known, and FadL may become saturated, inhibiting BTEX uptake (i.e., the added ^13^C-benzene in our experiment). Bacteria also exhibit varying preferences for BTEX substrates ^69^ and the benzenoid collection of hydrocarbons. Benzene may be of lower value to *Marinobacter* compared to the full collection of VOC substrates from growing *P. tricornutum,* or benzene may be oxidized to CO_2_ in the coculture rather than being incorporated into biomass as was observed in ASW and PTspent. Nevertheless, the contrasting ^13^C labeling results in *Marinobacter* suggest attachment dynamics in the *P. tricornutum* phycosphere have an important role in VOC uptake.

In the model set of phytoplankton and associated bacteria we investigated, phycosphere resources were exploited by bacteria employing varying strategies: VOC specialists (*Marinobacter* and *Roseibium*) that were motile and attached, generalists capable of VOC and macromolecular uptake (*Rhodophyticola* and *Stappia*), and macromolecular specialists (*Yoonia*). We hypothesize that VOC specialists use chemotaxis to find and colonize the phycosphere, where proximity to sources of low molecular weight, rapidly diffusing VOCs offers advantages ^28^. Unknown is the duration of attachment or how hydrocarbon uptake is differentially regulated in the presence and absence of *P. tricornutum*. The free-living generalists, *Rhodophyticola* and *Stappia*, used VOCs and carbohydrate DOC and have fewer genes encoding hydrocarbon oxidation proteins.

Collectively, VOC use by phycosphere bacteria was surprisingly broad in the range of compounds used and increased gross carbon production up to 29% in *P. tricornutum.* These findings add support to previous reports of VOC uptake by bacteria and stimulation of carbon fixation in phytoplankton cocultured with VOC oxidizers. Previous reports show that balanced production and consumption maintain low VOC concentrations in the oceans and have associated VOC consumption with specialized metabolism in abundant, streamlined, non-motile cells ^49,73–75^. On the contrary, in the set of bacteria we studied, some of the most active VOC utilizers were motile and attached to the phytoplankton. Our results suggest that VOC flux from algae can be intercepted by VOC specialists in the phycosphere, reducing ocean VOC accumulation and air-sea transfer and stimulating phytoplankton carbon production to replenish lost VOC resources. If this conceptual model is correct, then it is conceivable that the mechanisms we described result in a global increase in ocean photosynthetic carbon production and are an important conduit of direct transfer of GCP to bacteria that does not rely on the influence of protistan grazing and virus predation.

## Data availability

All data generated or analyzed during this study is included in this published article [and its supplementary information files].

## Acknowledgments

We thank Christina Ramon for lab assistance and Nicholas Baetge, Michael Sieler, and Chih Ping Lee for assistance with data analysis. This research was supported by award ID OCE1948163 to K.H.H. and S.J.G. from the National Science Foundation, Biological Oceanography Program. This research was supported in part by a fellowship to S.J.G. by the Hanse-Wissenschaftskolleg-Institute for Advance Study (HWK). Work at LLNL was Supported by the US Department of Energy’s (DOE) Genomic Science Program through the LLNL μBioSpheres Science Focus Area grant # SCW1039 and performed under the auspices of the DOE under contract DE-AC52-07NA27344.

## Author contributions

K.H.H., X.M., and V.G.P. conceived of the study. V.G.P. conducted all experiments with assistance from K.A., K.J., and L.C. and wrote the first manuscript draft with K.H.H. NanoSIMS was conducted by X.M. and P.K.W. Data analysis was done by V.G.P. All authors contributed to the final version of the manuscript.

## Ethics declarations

The authors declare no competing interests.

## References

1. Carlson, C. A., Liu, S., Stephens, B. M. & English, C. J. DOM production, removal, and transformation processes in marine systems. Biogeochemistry of Marine Dissolved Organic Matter 137–246 (2024) doi:10.1016/B978-0-443-13858-4.00013-7.

2. Halsey, K. H. & Giovannoni, S. J. Biological controls on marine volatile organic compound emissions: A balancing act at the sea-air interface. Earth Sci Rev 240, 104360 (2023).

3. Saha, M. & Fink, P. Algal volatiles – the overlooked chemical language of aquatic primary producers. Biological Reviews 97, 2162–2173 (2022).

4. Shemi, A., Ben-Dor, S., Rotkopf, R., Dym, O. & Vardi, A. Phylogeny and biogeography of the algal DMS-releasing enzyme in the global ocean. ISME Communications 3, (2023).

5. Xu, Q. et al. Volatile organic compounds released from *Microcystis flos-aquae* under nitrogen sources and their toxic effects on Chlorella vulgaris. Ecotoxicol Environ Saf 135, 191–200 (2017).

6. Zuo, Z. J., Zhu, Y. R., Bai, Y. L. & Wang, Y. Volatile communication between *Chlamydomonas reinhardtii* cells under salt stress. Biochem Syst Ecol 40, 19–24 (2012).

7. Dawson, R. A. et al. The microbiology of isoprene cycling in aquatic ecosystems. Aquatic Microbial Ecology 87, 79–98 (2021).

8. Asher, E., Dacey, J. W., Ianson, D., Peña, A. & Tortell, P. D. Concentrations and cycling of DMS, DMSP, and DMSO in coastal and offshore waters of the Subarctic Pacific during summer, 2010-2011. J Geophys Res Oceans 122, 3269–3286 (2017).

9. Pozzer, A. C., Gómez, P. A. & Weiss, J. Volatile organic compounds in aquatic ecosystems – Detection, origin, significance and applications. Science of the Total Environment 838, (2022).

10. Moore, E. R., Weaver, A. J., Davis, E. W., Giovannoni, S. J. & Halsey, K. H. Metabolism of key atmospheric volatile organic compounds by the marine heterotrophic bacterium *Pelagibacter* HTCC1062 (SAR11). Environ Microbiol 24, 212–222 (2022).

11. Rocco, M. et al. Oceanic phytoplankton are a potentially important source of benzenoids to the remote marine atmosphere. Communications Earth & Environment 2021 2:1 2, 1–8 (2021).

12. Misztal, P. K. et al. Atmospheric benzenoid emissions from plants rival those from fossil fuels. Scientific Reports 2015 5:1 5, 1–10 (2015).

13. Vila-Costa, M. et al. Phylogenetic identification and metabolism of marine dimethylsulfide-consuming bacteria. Environ Microbiol 8, 2189–2200 (2006).

14. Halsey, K. H., Padaki, V. G. & Giovannoni, S. The volatile organic carbon component of dissolved organic matter in the ocean. Biogeochemistry of Marine Dissolved Organic Matter 587–612 (2024) doi:10.1016/B978-0-443-13858-4.00001-0.

15. Moore, E. R., Davie-Martin, C. L., Giovannoni, S. J. & Halsey, K. H. *Pelagibacter* metabolism of diatom-derived volatile organic compounds imposes an energetic tax on photosynthetic carbon fixation. Environ Microbiol 22, 1720–1733 (2020).

16. Wu, H. jun et al. Metagenomic analysis reveals specific BTEX degrading microorganisms of a bacterial consortium. AMB Express 13, (2023).

17. Bacosa, H. P., Mabuhay-Omar, J. A., Balisco, R. A. T., Omar, D. M. & Inoue, C. Biodegradation of binary mixtures of octane with benzene, toluene, ethylbenzene or xylene (BTEX): insights on the potential of *Burkholderia, Pseudomonas* and *Cupriavidus* isolates. World J Microbiol Biotechnol 37, 1–8 (2021).

18. Hocinat, A., Boudemagh, A., Ali-Khodja, H. & Medjemadj, M. Aerobic degradation of BTEX compounds by *Streptomyces* species isolated from activated sludge and agricultural soils. Arch Microbiol 202, 2481–2492 (2020).

19. Khodaei, K., Nassery, H. R., Asadi, M. M., Mohammadzadeh, H. & Mahmoodlu, M. G. BTEX biodegradation in contaminated groundwater using a novel strain (*Pseudomonas* sp. BTEX-30). Int Biodeterior Biodegradation 116, 234–242 (2017).

20. Jiang, B. et al. Biodegradation of Benzene, Toluene, Ethylbenzene, and o-, m-, and p-Xylenes by the Newly Isolated Bacterium *Comamonas* sp. JB. Appl Biochem Biotechnol 176, 1700–1708 (2015).

21. Fu, H., Uchimiya, M., Gore, J. & Moran, M. A. Ecological drivers of bacterial community assembly in synthetic phycospheres. Proc Natl Acad Sci U S A 117, 3656–3662 (2020).

22. Seymour, J. R., Amin, S. A., Raina, J.-B. & Stocker, R. Zooming in on the phycosphere: the ecological interface for phytoplankton–bacteria relationships. Nat Microbiol 2, 17065 (2017).

23. Mitchell, J. G., Okubo, A. & Fuhrman, J. A. Microzones surrounding phytoplankton form the basis for a stratified marine microbial ecosystem. Nature 1985 316:6023 316, 58–59 (1985).

24. Seymour, J. R., Ahmed, T., Durham, W. M. & Stocker, R. Chemotactic response of marine bacteria to the extracellular products of *Synechococcus* and *Prochlorococcus*. Aquatic Microbial Ecology 59, 161–168 (2010).

25. Smriga, S., Fernandez, V. I., Mitchell, J. G. & Stocker, R. Chemotaxis toward phytoplankton drives organic matter partitioning among marine bacteria. Proc Natl Acad Sci U S A 113, 1576–1581 (2016).

26. Kogure, K., Simidu, U. & Taga, N. Bacterial attachment to phytoplankton in sea water. J Exp Mar Biol Ecol 56, 197–204 (1981).

27. Ramanan, R. et al. Phycosphere bacterial diversity in green algae reveals an apparent similarity across habitats. Algal Res 8, 140–144 (2015).

28. Raina, J. B. et al. Chemotaxis increases metabolic exchanges between marine picophytoplankton and heterotrophic bacteria. Nature Microbiology 2023 8:3 8, 510–521 (2023).

29. Ugolini, G. S. et al. Microfluidic approaches in microbial ecology. Lab Chip 24, 1394– 1418 (2024).

30. Durham, B. P. et al. Cryptic carbon and sulfur cycling between surface ocean plankton. Proc Natl Acad Sci U S A 112, 453–457 (2015).

31. Nouchi, I., Hosono, T. & Sasaki, K. Seasonal changes in fluxes of methane and volatile sulfur compounds from rice paddies and their concentrations in soil water. Plant Soil 195, 233–245 (1997).

32. Brisson, V., et al. Identification of Effector Metabolites Using Exometabolite Profiling of Diverse Microalgae. mSystems 6, (2021).

33. Russo, A., Pollastri, S., Ruocco, M., Monti, M. M. & Loreto, F. Volatile organic compounds in the interaction between plants and beneficial microorganisms. J Plant Interact 17, 840–852 (2022).

34. Yurimoto, H. & Sakai, Y. Interaction between C1-microorganisms and plants: contribution to the global carbon cycle and microbial survival strategies in the phyllosphere. Biosci Biotechnol Biochem 87, 1–6 (2022).

35. Samo, T. J. et al. Attachment between heterotrophic bacteria and microalgae influences symbiotic microscale interactions. Environ Microbiol 20, 4385–4400 (2018).

36. Di Costanzo, F., Di Dato, V. & Romano, G. Diatom–Bacteria Interactions in the Marine Environment: Complexity, Heterogeneity, and Potential for Biotechnological Applications. Microorganisms 2023, Vol. 11, Page 2967 11, 2967 (2023).

37. Wear, E. K., Carlson, C. A., Windecker, L. A. & Brzezinski, M. A. Roles of diatom nutrient stress and species identity in determining the short- and long-term bioavailability of diatom exudates to bacterioplankton. Mar Chem 177, 335–348 (2015).

38. Liu, S. et al. Opportunities and challenges of using metagenomic data to bring uncultured microbes into cultivation. Microbiome 10, (2022).

39. Wang, Y., Zhao, Y., Bollas, A., Wang, Y. & Au, K. F. Nanopore sequencing technology, bioinformatics and applications. Nature Biotechnology 2021 39:11 39, 1348–1365 (2021).

40. Van Tol, H. M., Amin, S. A. & Virginia Armbrust, E. Ubiquitous marine bacterium inhibits diatom cell division. ISME J 11, 31–42 (2017).

41. Helliwell, K. E., Shibl, A. A. & Amin, S. A. The Diatom Microbiome: New Perspectives for Diatom-Bacteria Symbioses. The Molecular Life of Diatoms 679–712 (2022) doi:10.1007/978-3-030-92499-7_23/COVER.

42. Balakrishnan, R., de Silva, R. T., Hwa, T. & Cremer, J. Suboptimal resource allocation in changing environments constrains response and growth in bacteria. Mol Syst Biol 17, 10597 (2021).

43. Nelson, D. M., Tréguer, P., Brzezinski, M. A., Leynaert, A. & Quéguiner, B. Production and dissolution of biogenic silica in the ocean: Revised global estimates, comparison with regional data and relationship to biogenic sedimentation. Global Biogeochem Cycles 9, 359–372 (1995).

44. Mayali, X. et al. Single-cell isotope tracing reveals functional guilds of bacteria associated with the diatom *Phaeodactylum tricornutum*. Nat Commun 14, 5642 (2023).

45. Kim, H. et al. Bacterial response to spatial gradients of algal-derived nutrients in a porous microplate. The ISME Journal 2021 16:4 16, 1036–1045 (2021).

46. Guillard, R. R. L. & Ryther, J. H. Studies of marine planktonic diatoms: *I. Cyclotella nana hustedt*, and *Detonula confervacea* (cleve) gran. Can J Microbiol 8, 229–239 (1962).

47. Ritchie, R. J. Consistent Sets of Spectrophotometric Chlorophyll Equations for Acetone, Methanol and Ethanol Solvents. Photosynth Res 89, 27–41 (2006).

48. Kolber, Z. S., Prášil, O. & Falkowski, P. G. Measurements of variable chlorophyll fluorescence using fast repetition rate techniques: defining methodology and experimental protocols. Biochimica et Biophysica Acta (BBA) - Bioenergetics 1367, 88–106 (1998).

49. Halsey, K. H. et al. Biological cycling of volatile organic carbon by phytoplankton and bacterioplankton. Limnol Oceanogr 62, 2650–2661 (2017).

50. Yáñez-Serrano, A. M. et al. GLOVOCS-Master compound assignment guide for proton transfer reaction mass spectrometry users. Atmos Environ 244, 117929 (2021).

51. Holzinger, R. et al. Validity and limitations of simple reaction kinetics to calculate concentrations of organic compounds from ion counts in PTR-MS. Atmos Meas Tech 12, 6193–6208 (2019).

52. Apweiler, R. et al. The Universal Protein Resource (UniProt) 2009. Nucleic Acids Res. 37, D169–D174 (2008).

53. Pearson, W. R. An Introduction to Sequence Similarity (“Homology”) Searching. Current protocols in bioinformatics / editoral board, Andreas D. Baxevanis … [et al.] **0 3**, 10.1002/0471250953.bi0301s42 (2013).

54. Edgar, R. C. MUSCLE: multiple sequence alignment with high accuracy and high throughput. Nucleic Acids Res 32, 1792 (2004).

55. Gu, Z. Complex heatmap visualization. iMeta 1, (2022).

56. Gu, Z., Eils, R. & Schlesner, M. Complex heatmaps reveal patterns and correlations in multidimensional genomic data. Bioinformatics 32, 2847–2849 (2016).

57. Lang, X. P., He, Z., Yang, G. P. & Dai, G. Physiological responses and altered halocarbon production in *Phaeodactylum tricornutum* after exposure to polystyrene microplastics. Ecotoxicol Environ Saf 268, 115702 (2023).

58. Cabrita, M. T., Vale, C. & Rauter, A. P. Halogenated Compounds from Marine Algae. Marine Drugs 2010, Vol. 8, Pages 2301-2317 8, 2301–2317 (2010).

59. Radman, S., Čagalj, M., Šimat, V. & Jerković, I. Seasonal Variability of Volatilome from *Dictyota dichotoma*. Molecules 2022, Vol. 27, Page 3012 27, 3012 (2022).

60. Jerković, I. et al. Phytochemical study of the headspace volatile organic compounds of fresh algae and seagrass from the Adriatic Sea (single point collection). PLoS One 13, (2018).

61. Koss, A. R. et al. Non-methane organic gas emissions from biomass burning: Identification, quantification, and emission factors from PTR-ToF during the FIREX 2016 laboratory experiment. Atmos Chem Phys 18, 3299–3319 (2018).

62. Jakoncic, J., Jouanneau, Y., Meyer, C. & Stojanoff, V. The catalytic pocket of the ring-hydroxylating dioxygenase from *Sphingomonas* CHY-1. (2006) doi:10.1016/j.bbrc.2006.11.117.

63. Tan, H. M., Tang, H. Y., Joannou, C. L., Abdel-Wahab, N. H. & Mason, J. R. The *Pseudomonas putida* ML2 plasmid-encoded genes for benzene dioxygenase are unusual in codon usage and low in G + C content. Gene 130, 33–39 (1993).

64. Bagnéris, C., Cammack, R. & Mason, J. R. Subtle Difference between Benzene and Toluene Dioxygenases of Pseudomonas putida. Appl Environ Microbiol 71, 1570 (2005).

65. Kotani, T., Yurimoto, H., Kato, N. & Sakai, Y. Novel acetone metabolism in a propane-utilizing bacterium, *Gordonia* sp. strain TY-5. J Bacteriol 189, 886–893 (2007).

66. Chen, Y. C., Peoples, O. P. & Walsh, C. T. *Acinetobacter* cyclohexanone monooxygenase: gene cloning and sequence determination. J Bacteriol 170, 781 (1988).

67. Mangwani, N., Kumari, S. & Das, S. Involvement of quorum sensing genes in biofilm development and degradation of polycyclic aromatic hydrocarbons by a marine bacterium *Pseudomonas aeruginosa* N6P6. Appl Microbiol Biotechnol 99, 10283–10297 (2015).

68. Olson, M. S., Ford, R. M., Smith, J. A. & Fernandez, E. J. Quantification of bacterial chemotaxis in porous media using magnetic resonance imaging. Environ Sci Technol 38, 3864–3870 (2004).

69. Lacal, J. et al. Bacterial chemotaxis towards aromatic hydrocarbons in *Pseudomonas*. Environ Microbiol 13, 1733–1744 (2011).

70. Matilla, M. A., Gavira, J. A. & Krell, T. Accessing nutrients as the primary benefit arising from chemotaxis. Curr Opin Microbiol 75, 102358 (2023).

71. Bell, W. & Mitchell, R. Chemotactic and growth responses of marine bacteria to algal extracellular products. Biol. Bull. 143, 265–277 (1972).

72. Hearn, E. M., Patel, D. R. & Van Den Berg, B. Outer-membrane transport of aromatic hydrocarbons as a first step in biodegradation. Proc Natl Acad Sci U S A 105, 8601–8606 (2008).

73. Sun, J. et al. One Carbon Metabolism in SAR11 Pelagic Marine Bacteria. PLoS One 6, e23973 (2011).

74. Simó, R., Cortés-Greus, P., Rodríguez-Ros, P. & Masdeu-Navarro, M. Substantial loss of isoprene in the surface ocean due to chemical and biological consumption. Communications Earth & Environment 2022 3:1 3, 1–8 (2022).

75. Davie-Martin, C. L., Giovannoni, S. J., Behrenfeld, M. J., Penta, W. B. & Halsey, K. H. Seasonal and Spatial Variability in the Biogenic Production and Consumption of Volatile Organic Compounds (VOCs) by Marine Plankton in the North Atlantic Ocean. Front Mar Sci 7, (2020).

