## Supplementary Fig. S1 for "Bacterial Volatile Organic Compound Specialists in the Phycosphere"

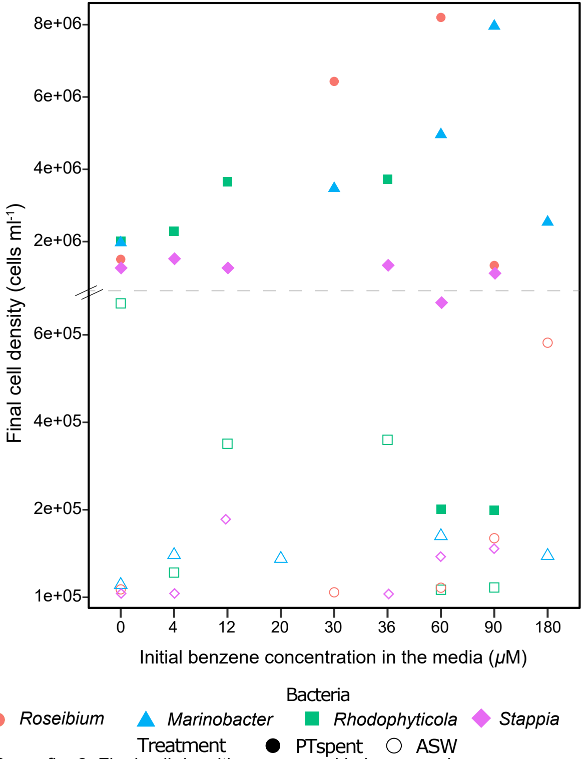


Supp. fig. 2: Final cell densities measured in benzene dose experiments. Each bacterium was inoculated into Ptspent (filled symbols) and ASW media (open symbols) with initial benzene concentrations 0-180 μM. *Roseibium* (circles), *Marinobacter* (triangles), and *Rhodophyticola* (squares).
