## Supplementary Fig. S2 for "Bacterial Volatile Organic Compound Specialists in the Phycosphere"

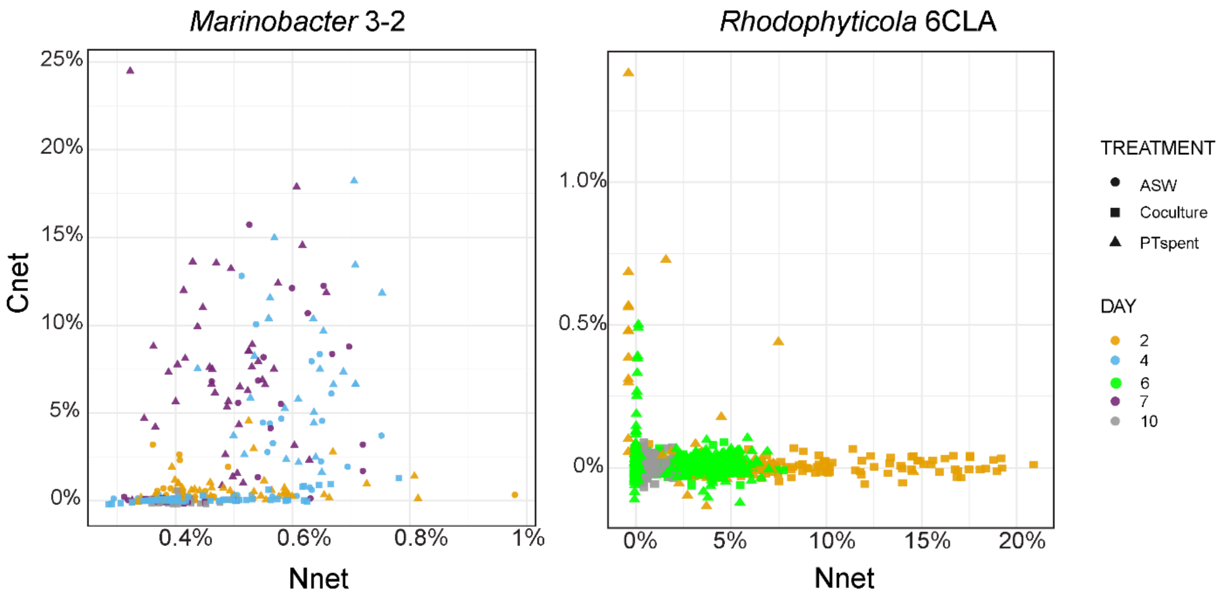


Supplementary Fig. S3: $C_{net}$ vs $N_{net}$ in *Marinobacter* and *Rhodophyticola*. Percent of bacterial biomass derived from $^{13}$C-benzene (Cnet) or $^{15}$N-leucine (Nnet) following incubation for two to ten days. $C_{net}$ and $N_{net}$ in killed cell controls are subtracted from $C_{net}$ and $N_{net}$ in growing *Marinobacter* and *Rhodophyticola.*

*NanoSIMS sample preparation*:

Samples (5 - 10 ml) were immediately fixed with 10% formalin, incubated for 30 min at 4° C, filtered onto 0.2 µm polycarbonate filters, and washed three times with nanopore Milli-Q water. To account for non-specific $^{13}$C binding, killed-controls were grown with no benzene, fixed with formaldehyde, incubated with $^{13}$C benzene for 30 min at 4° C, filtered, and washed. All filters were air-dried and then cut into small wedges, adhered to aluminum disks using conductive tabs (#16084-6, Ted Pella, Redding, CA), and shipped at ambient temperature to Lawrence Livermore National Laboratory. PT-*Rhodophyticola* required a hydrofluoric acid cleaning (1% HF for 10 min followed by rinse in Milli-Q water) due to a coating of particulate matter blocking cells.

*Brief explanation of isotopic measurement using nanoSIMS*:

Samples were sputter-coated with ~5 nm of gold. Analysis locations were first sputtered to a depth of ~60 nm with the primary $^{133}Cs^{+}$ ion beam set to ~100 pA before analysis with 2 pA (150 nm beam diameter at 16 keV). Rastering was performed over 20 x 20 µm analysis areas with a dwell time of 1 ms pixel$^{-1}$ for 19-30 scans (cycles) and generated images containing 256 x 256 pixels, with sputtering equilibrium to a depth of ~60 nm. After tuning the secondary ion mass spectrometer for mass resolving power of ~7000 (1.5x corrected), secondary electron images and quantitative secondary ion images were simultaneously collected for $^{12}C_2^{-}$, $^{13}C^{12}C^{-}$, $^{12}C^{14}N^{-}$, and $^{12}C^{15}N^{-}$ on individual electron multipliers in pulse counting mode. All NanoSIMS datasets were initially processed using L'Image (http://limagesoftware.net) to perform dead time and image shift correction of ion image data before creating $^{13}C/^{12}C$ and $^{15}N/^{14}N$ ratio images. To quantify substrate incorporation, cells were identified based on $^{12}C^{14}N^{-}$ images, and regions of interest (ROIs) were drawn manually around each bacterial cell. In order to calculate how much C or N a cell incorporated from a substrate that was isotopically labeled, we calculated $X_{net}$, either $C_{net}$ or $N_{net}$ (Dekas et al., 2019; Pett-Ridge et al., 2022), which represent the fraction of a cell's biomass that originated from the added isotope labeled substrate (here, $^{13}C$-benzene and $^{15}N$-leucine for $C_{net}$ and $N_{net}$, respectively) and used the killed control ($f_{X_i}$), the measurement at the final timepoint ($f_{X_f}$), and the isotope fraction of the substrate ($f_{X_s}$), which assumed negligible unlabeled benzene ($^{12}C$-benzene):

$$X_{net\%} = \frac{f_{X_f} - f_{X_i}}{f_{X_s} - f_{X_i}} \times 100\%$$
