## Supplementary Fig. S3 for "Bacterial Volatile Organic Compound Specialists in the Phycosphere"

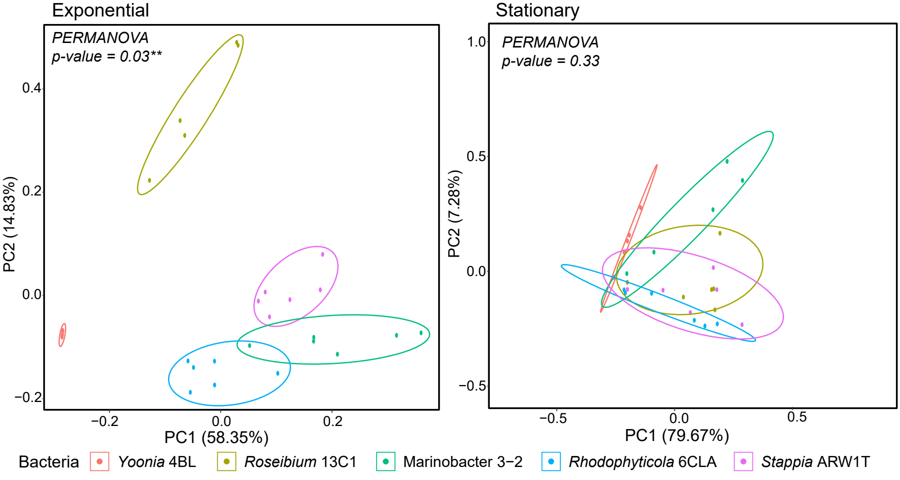


Supp. fig. 1: Principal coordinate analysis of depleted VOCs in *P. tricornutum*–bacteria cocultures in exponential (left) and stationary (right) growth phases.
